# What lurks in the dark? - An innovative framework for studying diverse wild insect microbiota

**DOI:** 10.1101/2024.08.26.609658

**Authors:** Karol H. Nowak, Emily Hartop, Monika Prus-Frankowska, Mateusz Buczek, Michał Kolasa, Tomas Roslin, Otso T. Ovaskainen, Piotr Łukasik

**Author notes:** The datasets supporting the conclusions of this article are available in the GitHub repository, https://github.com/KarolHub/Phorid-Microbiota.

## Abstract

Symbiotic microorganisms can profoundly impact insects, their life history traits, population dynamics, and evolutionary trajectories. However, microbiota remain poorly understood in natural insect communities, especially in ‘dark taxa’ - i.e., hyperdiverse, yet understudied clades.

Here, we implemented a novel multi-target amplicon sequencing approach to study microbiota in complex, species-rich communities. It combines four methodological innovations: (1) To establish a host taxonomic framework, we sequenced amplicons of the host marker gene (COI) and reconstructed barcodes alongside microbiota characterisation. (2) To assess microbiota abundance, we incorporated spike-in-based quantification. (3) To improve the phylogenetic resolution for the dominant endosymbiont, *Wolbachia*, we analysed bycatch data from the COI amplicon sequencing. (4) To investigate the primary drivers of host-microbe associations in massive multi-dimensional datasets, we performed HMSC modelling.

Applying this approach to 1,842 wild-caught scuttle flies (Diptera: Phoridae) from northern Sweden, we organised them into 480 genotypes and 186 species and gained unprecedented insights into their microbiota. We found orders-of-magnitude differences in bacterial abundance and massive within-population variation in microbiota composition. Patterns and drivers differed among microbial functional categories: the distribution and abundance of facultative endosymbionts (*Wolbachia*,

*Rickettsia*, *Spiroplasma*) were shaped by host species, genotype and sex. In contrast, many other bacterial taxa were broadly distributed across species and sites.

This study highlights facultative endosymbionts as key players in insect microbiota and reveals striking variations in distributional patterns of microbial clades. It also demonstrates the power of integrative sequencing approaches in uncovering the ecological complexity and significance of symbiotic microorganisms in multi-species natural communities.

## 1. Introduction

Microbial symbioses have played tremendously important roles in the biology and evolution of higher organisms (1). The many levels at which different functional categories of symbionts have shaped hosts’ biology are relatively well-understood in insects (2). The range of microbial symbionts’ effects on insect life-history traits - nutrition, reproduction, and defence (3,4) translate to population processes and patterns (5–7), host evolutionary trajectories (8,9), and effects at the level of multi-species communities (10). There is little doubt that through such combined effects, microbiota have shaped the global biodiversity and will continue to play key roles in ecosystems rapidly changing under anthropogenic pressures (11,12).

Despite their massive importance, insect microbiota are poorly known outside of model species. Some biological systems including honey bees (13) or pea aphids (14) have been the focus of extensive research on symbiosis diversity, function, mechanisms, and various other aspects. However, for some 6 million of the remaining insect species on our planet (15,16), our understanding of microbiota is nonexistent, or at best, very limited. Among the primary reasons is the sheer diversity of insects and microorganisms, which poses a significant challenge in generating and handling vast amounts of multidimensional ecological data. We lack reliable and cost-effective workflows for processing large numbers of wild-caught insects, encompassing host identity validation and microbiome characterisation, while including controls for different types of contamination (17–19). Additionally, existing approaches rarely address questions about microbial abundance. Yet, such variation is critical, as insects differ dramatically in microbial load (20). Data analysis is another overlooked area with challenges arising when microbial communities are dominated by one or a few strains, when overall microbial abundance is low, and when contamination is not controlled for. Finally, we lack adequate analytical solutions for the reconstruction of microbiome patterns and drivers in highly multidimensional datasets.

As a result of these challenges, we know very little about the microbial composition and abundance across the arthropod diversity – or the drivers of this variation. Host biology, especially the diet, has been identified as an important driver (8), alongside host taxonomy (21). However, even within species, there can be major differences among populations, within a population over time, and among individuals sampled at the same time from a single population (22). Similarly, the differences in microbiota between host sexes have rarely been considered outside of the context of reproductive manipulation (23,24). Interactions among these and other possible microbiota drivers can make analyses especially challenging. At the same time, we would anticipate that microbial taxa and functional categories would differ in how the infections are determined. A systematic assessment of these and other host-microbe interaction drivers would require a combination of comprehensive laboratory methods applied at a large scale and robust data analysis approaches.

Much of the global insect diversity is made up of so-called ‘dark taxa’: insect clades that are hyperdiverse, abundant, and broadly distributed, yet remain poorly studied (25). The study of dark taxa has greatly benefitted from recent advances in molecular workflows (26,27), particularly in Megabarcoding - high-throughput barcoding of single specimens (28), which has enabled large-scale analyses of hyperdiverse insects on a broader scale than previously possible (29). Growing barcoding data provide a promising opportunity to illuminate the hidden microbial diversity across these enigmatic clades.

Scuttle flies (Diptera: Phoridae) are a quintessential example of a ‘dark taxon’: they are hyperdiverse, abundant, and globally distributed, but remain poorly understood (25,30). They exhibit diverse life strategies, including parasitism, predation, herbivory, and saprophagy (31). Such varied life histories allow them to occupy numerous ecological niches, from bustling urban centres (32) and subterranean caves (33) to intertidal zones (34) and the cavities of pitcher plants (35). Despite their ubiquity and diverse ecological roles, our understanding of their diversity, distribution, and biology remains grossly incomplete (26). Given our poor state of knowledge about scuttle flies generally, it is not surprising that little is known about their interactions with microbial symbionts. Only a few studies have addressed questions about phorid-associated microbiota, using one or few species and with shallow sampling depth (36,37).

In this study, we leverage a novel combination of molecular and analytical approaches to comprehensively characterise microbiota abundance, composition, and distribution in diverse scuttle flies across six sites in Northern Sweden. To resolve variation in microbiota composition among scuttle flies, we employ high-resolution COI amplicon sequencing for host identification, bacterial 16SV4 rRNA sequencing using synthetic spike-ins to quantify microbial abundance, and state-of-the-art Hierarchical Modelling of Species Communities (HMSC) to disentangle ecological drivers of microbial variation. By integrating these innovative techniques, we assess how microbial communities are shaped by site, season, host species identity, and individual host traits such as sex and genotype. This innovative methodological pipeline provides an unprecedented resolution of host-microbiota interactions in a hyperdiverse insect group, advancing our understanding of the ecological and evolutionary forces shaping microbial associations in wild communities.

## 2. Materials and methods

### 2.1. Field collections

Samples were collected using Malaise traps as a part of the Swedish Insect Inventory Project (SIIP) (38). The collection was carried out in northern Sweden at six sites, separated by distances varying from less than 50 m to about 390 km (for exact locations, see Supplementary Table S1 and Fig. 1). Selected samples come from late spring (June to July) and summer (July to August) of 2018 (Supplementary Table S1). After collection, samples in 95% ethanol were stored at −20 °C in Station Linné, Sweden, where taxonomists manually sorted the scuttle flies from the samples. We randomly selected 88 males and 88 females for processing from each of the six sites and two seasons (except for three combinations where only 87, 86 and 84 individuals were available; see Supplementary Table S1 for details), totalling 2,105 insects.

**Fig. 1:**
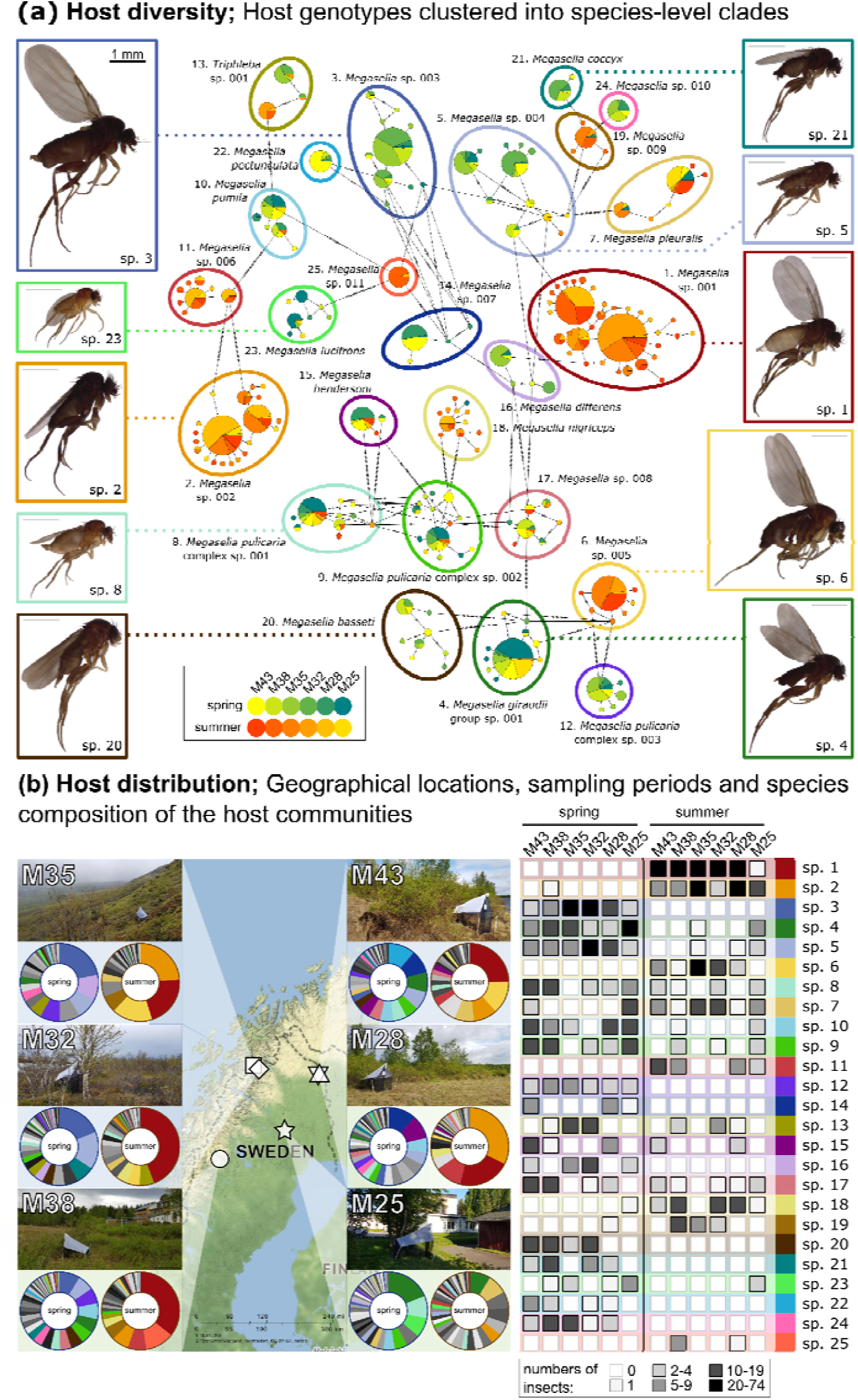
(**a**) Haplotype network of the 25 most abundant scuttle fly species highlights the diversit among the sampled insects, and showcases challenges with their delimitation. Individual pie charts represent fly COI genotypes, coloured by collection site and season. Selected species are represented by photos of individually barcoded individuals from various sites, with all images scaled uniformly for consistency. Genotypes of the remaining 161 species are not shown. The network was drawn using PopArt (58). (**b**) The distribution and diversity of the 25 most commonly sampled scuttle fly species from Northern Sweden highlights community composition variation among the two seasons and overlap among the six sampled sites.

### 2.2 DNA extraction, amplicon library preparation and sequencing

The insects were processed and prepared for sequencing in the Molecular Ecology Laboratory at the Institute of Environmental Sciences of Jagiellonian University, following a protocol comprehensively described recently (39). First, each insect was placed in a 2 ml tube with 190 µl of the lysis buffer (0.4 M NaCl, 10 mM Tris-HCl, 2 mM EDTA, 2% SDS), 10 µl proteinase K and different-sized ceramic beads. After two rounds of 30-second homogenisation on a homogenizer (Bead Ruptor Elite, Omni International), the samples were incubated for 2 hours at 55 °C. Subsequently, 40 µl aliquot of the homogenate was combined with 2 µl of the solution containing 10,000 or 20,000 copies (depending on the batch) of plasmids carrying artificial 16S rRNA amplification targets, Ec5502 (40), placed in a 96-well PCR plate, and purified using SPRI beads (41). DNA was eluted in 20 µl of TE buffer.

The amplicon libraries were prepared using a two-step PCR procedure (see method 4 (42)). In the first step, two marker regions of interest were simultaneously amplified using QIAGEN Multiplex Master Mix and template-specific primers with variable-length inserts and Illumina adapter stubs. Specifically, for the V4 region of bacterial 16S rRNA, we used primers 515F: GTGYCAGCMGCCGCGGTAA and 806R: GGACTACNVGGGTWTCTAAT (43), and for a 418 bp portion of the mitochondrial cytochrome oxidase I (COI) gene, primers BF3: CCHGAYATRGCHTTYCCHCG and BR2: TCDGGRTGNCCRAARAAYCA (44). The PCR reaction mix also contained 500 copies of another plasmid carrying an artificial 16S rRNA target, Ec5001 (40), but it was not used in this study and was subsequently filtered out from the data. SPRI bead-purified first-step PCR products were used as a template for the second indexing PCR. The libraries were verified on an agarose gel and based on band intensities pooled approximately equimolarly. The purified pool was quality checked using BioAnalyzer 2100 (Agilent), combined with pools representing other projects, and submitted to Swedish National Genomics Infrastructure for sequencing on Illumina NovaSeq SPrime flow cells (2×250 bp read mode). To control for contamination from different sources, for each of the 88 biological samples processed, we used eight different controls: negative controls for the DNA extraction (3), first PCR (2), and indexing PCR stages (1), as well as positive controls (2). Raw sequence data for all samples that passed filtering criteria and controls are available under GenBank BioProject accession number PRJNA1099311.

### 2.3. Bioinformatic analysis of amplicon data

The amplicon data were processed using scripts and files available at the GitHub repository, together with supplementary data: https://github.com/KarolHub/Phorid-Microbiota.

#### 2.3.1. Basic read sorting and processing

Initially, we split the demultiplexed reads into batches corresponding to COI and 16S-V4 targets using a custom Python script (Multisplit.py), which divided the files based on primer sequences and trimmed the primers. Further steps relied on the custom LSD.py script, a wrapper tool executing several steps using PEAR (45), VSEARCH (46), and USEARCH (47) programs. Specifically, we merged forward and reverse reads into contigs, discording those outside of the expected length range (400-430 for COI and 250-260 bp for 16S-V4), or with lower quality regions (Phred Score ≤ 30). Datasets representing both regions were then dereplicated to retain unique sequences per sample and their occurrence counts, with singleton genotypes discarded. We denoised each library separately using the UNOISE algorithm (48). Then, we clustered the zero-radius operational taxonomic units (ZOTUs) into operational taxonomic units (OTUs) using a 97% identity cutoff. Finally, we assigned taxonomy using the SINTAX algorithm (49), and taxonomic reference databases: MIDORI version GB 239 (50), with alphaproteobacterial sequences added, for the COI data and SILVA (51) version 138 SSU for 16S rRNA data. We retained identification to the lowest level that had at least an 80% classification confidence score.

#### 2.3.2. COI data analysis

The COI data contained a mix of sequences of different origins, including those representing insect mitochondrial genes, nuclear pseudogenes, alphaproteobacterial symbionts, and others. To obtain insect barcodes, we discarded sequences shorter or longer than the anticipated 418 nucleotides, as well as bacterial, non-arthropod, unassigned, and chimeric sequences. Next, we assessed the number of reads in the most abundant OTU classified as an arthropod and calculated its ratio relative to all Arthropoda OTUs combined. We excluded libraries from further analyses if they had fewer than 300 reads of the dominant OTU or if the aforementioned read ratio was less than 95%. Then, we designated the most abundant sequence within the dominant OTU of each sample as the individual’s barcode, dereplicated these barcodes, and clustered them with sequences from earlier barcoding studies (25) to generate a dendrogram. Given the complex taxonomy of phorid flies, we manually reviewed the proximity of barcodes and sequences of established phorid species within the dendrogram to determine which groups of barcodes form distinct species-level clades. Once we had the sequences divided into species-level clusters of barcodes, we compared them with sequences of named species from the same study (25), to obtain the taxonomy labels. The species names were assigned if a sequence matched the reference by at least 97%. Variants with no close matches were compared against the NCBI nt database using the BLAST algorithm, retaining labels with a match of 97% or higher.

The COI data contained sequences assigned to the order Rickettsiales, which we separated for further analyses. We filtered out non-*Wolbachia* sequences, chimeric sequences, and those of length other than 418 bp. This resulted in tables on *Wolbachia* COI ZOTU and OTU read number distribution among samples, analogous to standard bacterial 16S rRNA-based tables.

#### 2.3.3. 16S rRNA amplicon data filtering and abundance estimation

We decontaminated the 16S-V4 data from 1,845 samples with reliable insect COI barcodes using a custom QUACK.py script. This script classified as contaminants and removed ZOTUs unless the percentage of reads representing that ZOTU in at least one experimental sample was at least five times greater than the maximum percentage in any of the blank samples (52,53). It also removed sequences classified as Archaea, Eukaryota, chloroplasts, mitochondria and chimaeras. After calculating the ratio between the spike-in and remaining symbiont reads in each sample, the spike-in sequences were also removed. Afterwards, we calculated the absolute abundance of non-contaminant bacterial 16S rRNA copies in each sample by multiplying the symbiont to extraction spike-in read number ratio, the number of inserted extraction spike-in copies (10,000 or 20,000), and the inverse proportion of insect homogenate used during the DNA extraction (5). The abundance was estimated separately for each bacterial ZOTU, with the total bacterial abundance in the sample estimated as the summed abundance of individual ZOTUs. Three samples without extraction spike-in reads were discarded, along with control samples, resulting in the final total number of phorid samples of 1842. For subsequent comparisons and modelling, we used tables on bacterial OTU and ZOTU estimated 16S rRNA copy number distributions across samples.

#### 2.3.4. Analyses

To examine the drivers of variation in insect microbiomes we used Hierarchical Modelling of Species Communities (HMSC; (54,55), a type of joint species distribution modelling (JSDM; (56)). Focusing on 75 species represented by at least 5 individuals (for a total of 1634 samples), we quantified how ecological and host-related factors shape microbial community composition and abundance. To achieve this, we fitted separate HMSC models for each of three response variables: (1) estimated copy numbers for the 50 most abundant 16S-V4 OTUs, (2) estimated copy numbers for the 100 most dominant 16S-V4 ZOTUs, and (3) read counts of the 24 most dominant Wolbachia COI ZOTUs. For covariates, we used effects of season (spring vs. summer), site (n = 6), sex of the host (male vs. female), host species (n = 75), host genotype (n = 328), and sampling unit (n = 1,634).

Because our data were zero-inflated—with many taxa being absent from individual samples, resulting in numerous zero counts—we used a hurdle modelling approach to separately analyse presence-absence and abundance conditional on presence. First, we modelled the presence-absence of each taxon using a Bernoulli distribution with a probit link function. Then, for taxa that were present, we modelled abundance by log-transforming and scaling the response, assuming a normal distribution. In our models, season and sex were included as fixed effects, while site, host species, host genotype, and individual host insect were treated as community-level random effects, structured using latent variables to account for shared variation across taxa.

We fitted the model using the Markov Chain Monte Carlo (MCMC) approach implemented in the R-package HMSC (57) with default prior distributions (55). The posterior distribution was sampled with four chains, with 375 * thin MCMC iterations per chain. We ignored the first 125 * thin iterations as burn-in and thinned the remaining iterations by thin to yield 250 samples per chain and hence 1000 samples in total. The thinning parameter was increased as thin = 1, 10, 100,…until satisfactory MCMC convergence was reached. We considered MCMC convergence satisfactory if the third quartile of the potential scale reduction factors over the parameters measuring the responses of the taxa to season and sex was at most 1.05; this was reached with thin = 1000 for 16S-V4 ZOTU and *Wolbachia* COI ZOTU presence-absence models, and with thin = 100 for the other four model components.

To evaluate model fit, we focused on explanatory and predictive powers – the former based on fitting the models to all data and the latter through two-fold cross-validation. As metrics for evaluation, we used AUC for presence-absence models and R^2^ for models on abundance conditional on presence. To examine the importance of the fixed and random effects in structuring microbiome variation, we applied a variance partitioning approach to the explained variation. In other words, we examined what proportion of the overall variation explained by the model could be attributed to the individual factors and covariates. To examine the effects of sex and season, we counted the proportion of taxa that showed either a positive or negative response with at least 95% posterior probability. To examine patterns of co-variation not explained by sex or season, we examined which pairs of bacterial taxa showed a positive or negative association for each random effect (site, species, genotype and individual) with at least 90% posterior probability. The latter approach will reveal what microbial taxa appear to covary in their presence and abundance beyond what can be expected based on the fixed effects of sex and season.

## 3. Results

After stringent quality filtering of 2,105 amplicon sequencing libraries for COI and 16S-V4 marker regions of individual phorid flies, we retained on average 77 individuals for each of the 24 site-season-sex combinations - 936 males and 906 females, or 1,842 individuals in total (Supplementary Table S1). For individuals passing the filtering criteria, we obtained, on average, 27.1K (K = thousand) COI reads (min 373, max 191.9K, median 20.4K) (Supplementary File S1). We then used these data to reconstruct insect barcodes and classify specimens to species (Supplementary Table S2; see section 3.1), but also to reconstruct and compare COI sequences of *Wolbachia* (see section 3.3). We obtained, on average, 29.7K 16S-V4 reads (range 682 - 295.7K, median 19.5K) per experimental library. After filtering and removal of putative laboratory and reagent-derived contaminants and removing spike-in control reads, we retained, on average, 15.2K 16S-V4 non-contaminant reads (range 29 - 227K, median 5.4K) (Supplementary Table S3). We used these data to reconstruct microbial abundance and diversity across individuals (section 3.2).

### 3.1. Abundance, diversity and distribution of phorid flies

Of the COI amplicon data for individual phorid flies, ca. 99.5% of reads were classified as matching insects. On average, the most abundant insect ZOTU in each library, designated as the individual’s barcode, accounted for 98.8% of the filtered insect reads (min 52.2%, max 100%, median 99.5%). The COI barcodes of 1,842 phorid individuals represented 480 genotypes. Through a comparison with previously sequenced and morphologically characterised material, these genotypes we grouped into 186 species (Supplementary Table S2) (25). The number of clustered genotypes and their abundance varied among species (Fig. 1a). For example, the five most abundant species were represented by between 10 and 33 genotypes each (85 genotypes in total), but 59 of these genotypes were represented by a single individual only. Many of these low-abundance genotypes differed from the more abundant genotypes by a single nucleotide. 142 of the COI genotypes, representing 438 individuals, were confidently assigned to 63 recognised, named species, mostly within the genus *Megaselia*. A further 42 genotypes were assigned to two species complexes: the *Megaselia pulicaria* complex (132 individuals) and the *Megaselia giraudii* complex (82 individuals). Most of the samples (295 genotypes, 1176 individuals) were assigned to the genus level, and one genotype consisting of 14 individuals was assigned to the family level only. Given the low prevalence of individual genotypes, there was little indication of geographic structuring within species (Fig. 1, Supplementary Table S2). To allow unequivocal reference to each species (whether recognised or not), we assigned unique numbers to all species, generally corresponding to their rank abundance (nonetheless, subsequent changes to the sample set caused some discrepancies between rank and number.) We are referring to species by these numbers throughout the text (e.g., “sp. 1” and “sp. 15” - rather than “*Megaselia* sp. 001” and “*Megaselia hendersoni”*) (Fig. 1a, Supplementary Table S2).

Species’ representation in the dataset was highly skewed, with a few abundant species and a long tail of rare ones (Supplementary Table S2). Within the same season, we found an overlapping set of abundant species across all sites (Fig. 1b). For example, the five most abundant species were found at all six sites. At the same time, there was little overlap in community composition among seasons, with most species well-represented either in spring or in the summer (Fig. 1b). NMDS based on Bray-Curtis distances visibly separated spring and summer community samples (Supplementary Fig. S1).

### 3.2. Abundance and composition of phorid microbiota

Across 1,842 phorid samples, a total of 28M (M = million) reads scored as non-contaminants and represented 58.1K ZOTUs that clustered into 9.6K 97% OTUs (Supplementary File S3). Using the ratio of quantification spike-in to non-contaminant reads as an estimate of the abundance of specific bacterial ZOTUs, OTUs, and total bacteria (39,40), we were able to assess and interpret microbial abundance and diversity at the same time.

In terms of microbe abundance, we observed high variation in the symbiont to spike-in ratios across the samples. The ratio ranged between 0.015 and 354.8 (median 1.38), and with 10K or 20K artificial template spike-in copies added to one-fifth of insect homogenate before purification, these ratios translate to bacterial 16S rRNA copy numbers per insect ranging between 766 and 24M (median 815.4K) (Supplementary File S3). Even among individuals representing the same sex and species, the range in 16S rRNA copy numbers typically spanned between two to four orders of magnitude. At the same time, microbial abundance was often higher in females than males of the same species, by about an order of magnitude on average (Fig. 2a).

**Fig. 2:**
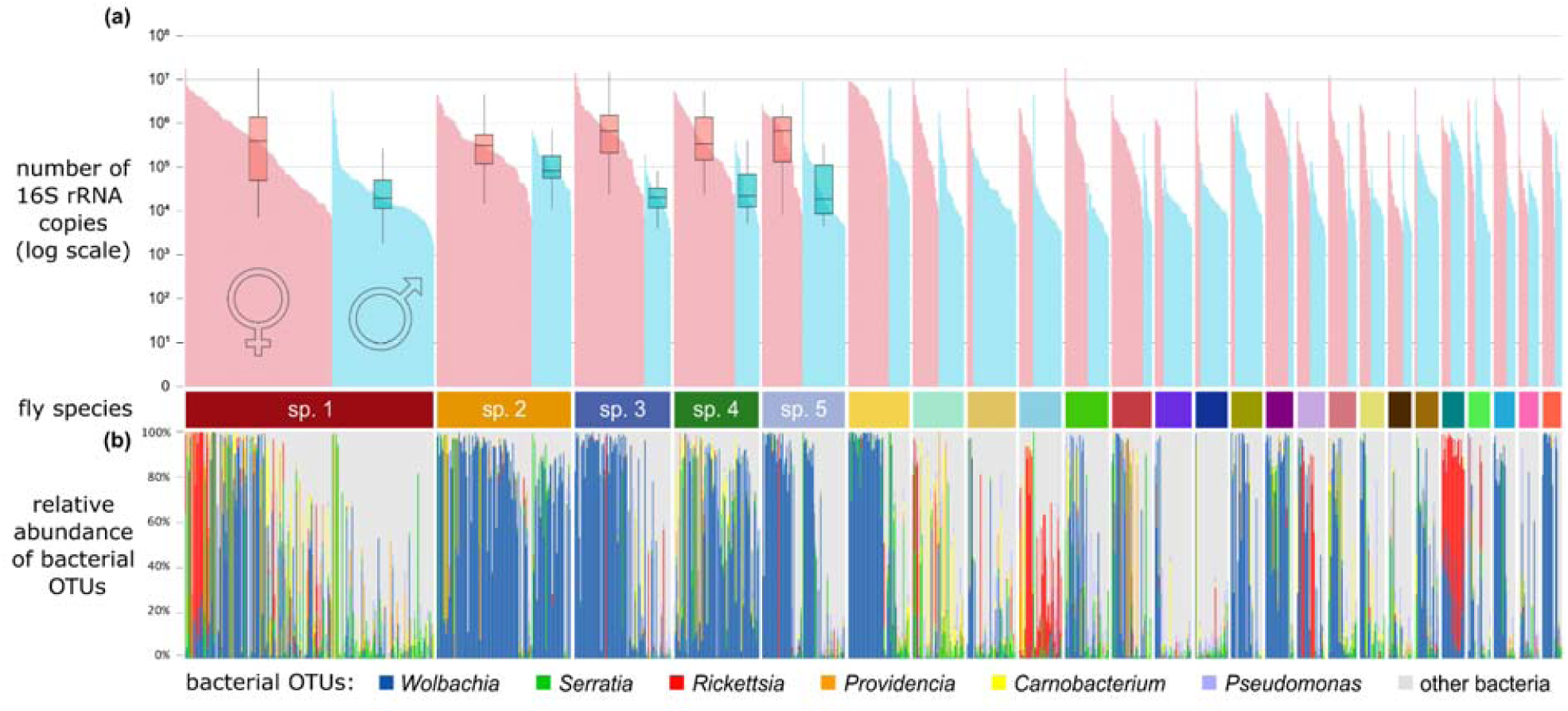
Simultaneous inspection of bacterial absolute abundance and microbial diversity data indicate strong variation across 1,193 specimens of the 25 dominant scuttle fly species. (**a**) The distribution of bacterial 16S-V4 rRNA copy numbers across individual insects, estimated based on quantification spike-ins added during the DNA extraction step. Here, each fly is represented by a narrow vertical bar, with bars sorted first by species assignment, then by sex, and then by abundance. (**b**) The relative abundances of the six most abundant bacterial 97% OTUs across individual insects were sorted as in panel **a**. The category ‘other bacteria’ shows all the remaining bacterial OTUs combined.

Although the number of detected 16S-V4 97% OTUs was high, phorid fly communities were dominated by relatively few bacterial clades (Supplementary Table S4). The most common OTU1, taxonomically assigned to the genus *Wolbachia*, was represented by 30.3% of decontaminated bacterial reads in a library. The next-most prevalent symbiont OTUs, *Serratia*, *Rickettsia*, *Providencia*, *Carnobacterium,* and *Pseudomonas,* comprised an average of 5.5%, 5.1%, 3.0%, 2.9%, and 2.7% of reads in a library, respectively. We identified multiple additional, less abundant bacterial OTUs representing the same six genera, plus a long list of other genera (Supplementary Table S4). These OTUs were broadly distributed across species, but their abundances showed non-random distribution patterns among individuals within species (Table 1; Fig. 2b). For example, *Rickettsia* comprised a particularly large share of the microbiota of females of species 1 with high overall bacterial abundance.

The analysis of abundance and distribution of dominant bacterial clades revealed large variation among individuals, even of the same species, population, and sex - as well as the complex interactions among likely drivers of microbial abundance and composition (Fig. 3). For example, when present, *Rickettsia* can be a quantitatively dominant microbe in females of phorid sp. 1, but its prevalence varies widely among populations. At a specific site (M38), *Providencia* is common and abundant in both sexes of phorid sp. 1 and sp. 2, but it is much less common or abundant at other sites, or in co-occurring phorid sp. 3.

**Fig. 3:**
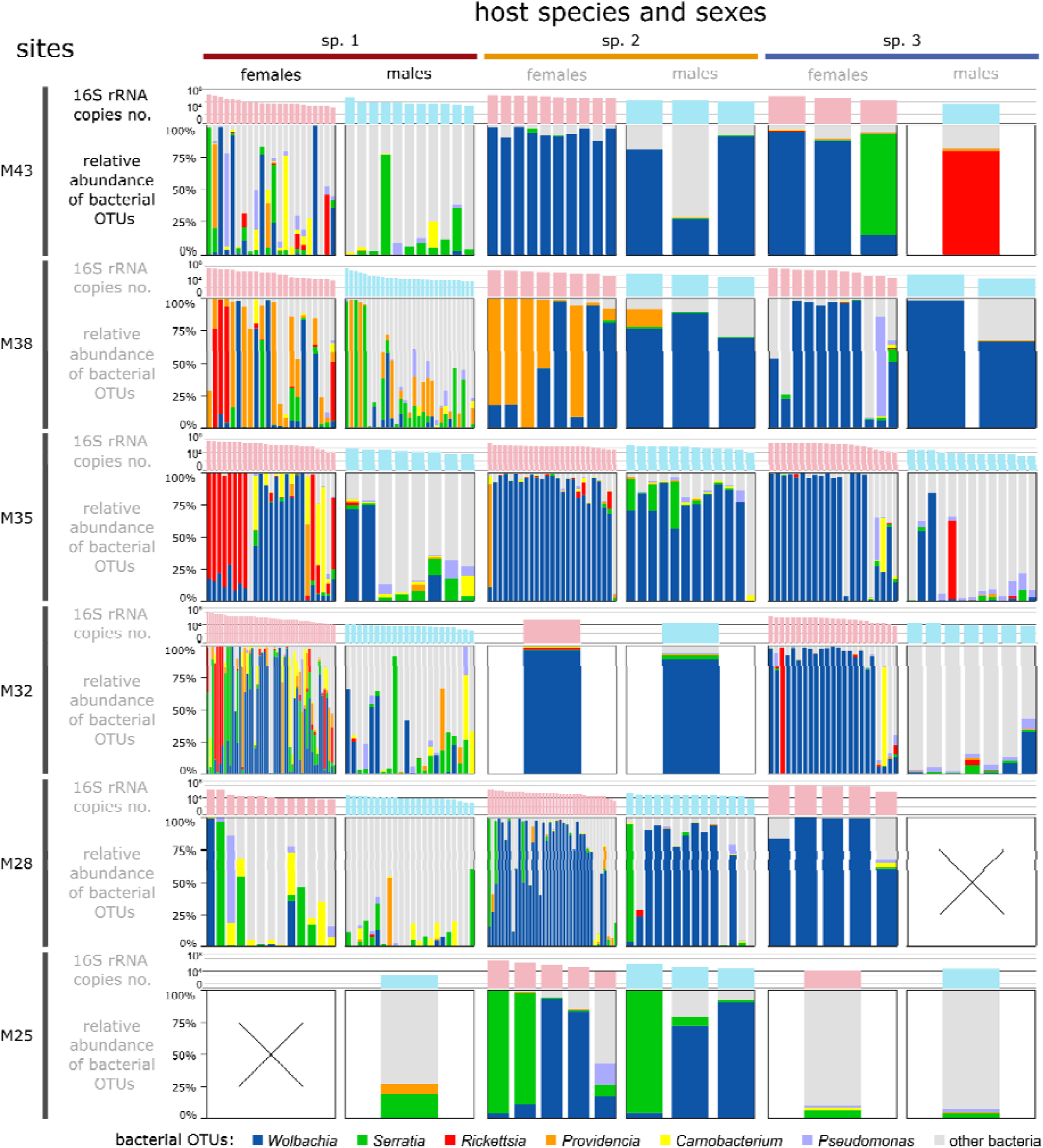
The comparison of total bacterial abundance and relative abundance of dominant bacterial OTUs showcased the extent and patterns of variation among the three most abundant phorid species, the two sexes, and the six sites. Each panel represents a particular species-sex-site combination, with the top bars indicating bacterial abundance, and the lower stacked bars, the relative abundance of th most abundant bacterial 16S OTUs (as in Fig. 2). Individual columns of varying width represent individual flies.

Inspection of the data at higher phylogenetic resolution - 16S rRNA ZOTUs instead of 97% OTUs - provided additional information about the host-microbe association patterns. Bacterial OTUs typically comprise several relatively abundant genotypes (ZOTUs), and these OTUs differ in how their component ZOTUs are distributed across host species (Fig. 4). Specifically, insect species were found to preferentially associate with only one or two specific 16S rRNA genotypes of facultative endosymbiont *Wolbachia* and *Rickettsia*. Different species associate with different facultative endosymbiont genotypes, and conversely, different facultative endosymbiont genotypes colonise different sets of species. In contrast, for most other abundant bacterial OTUs including *Serratia*, *Providencia*, *Carnobacterium*, and *Pseudomonas*, associations appear much less specific. Most insect species harbour several genotypes from these OTUs, and genotypes are broadly distributed across host species (Supplementary File S3).

**Fig. 4:**
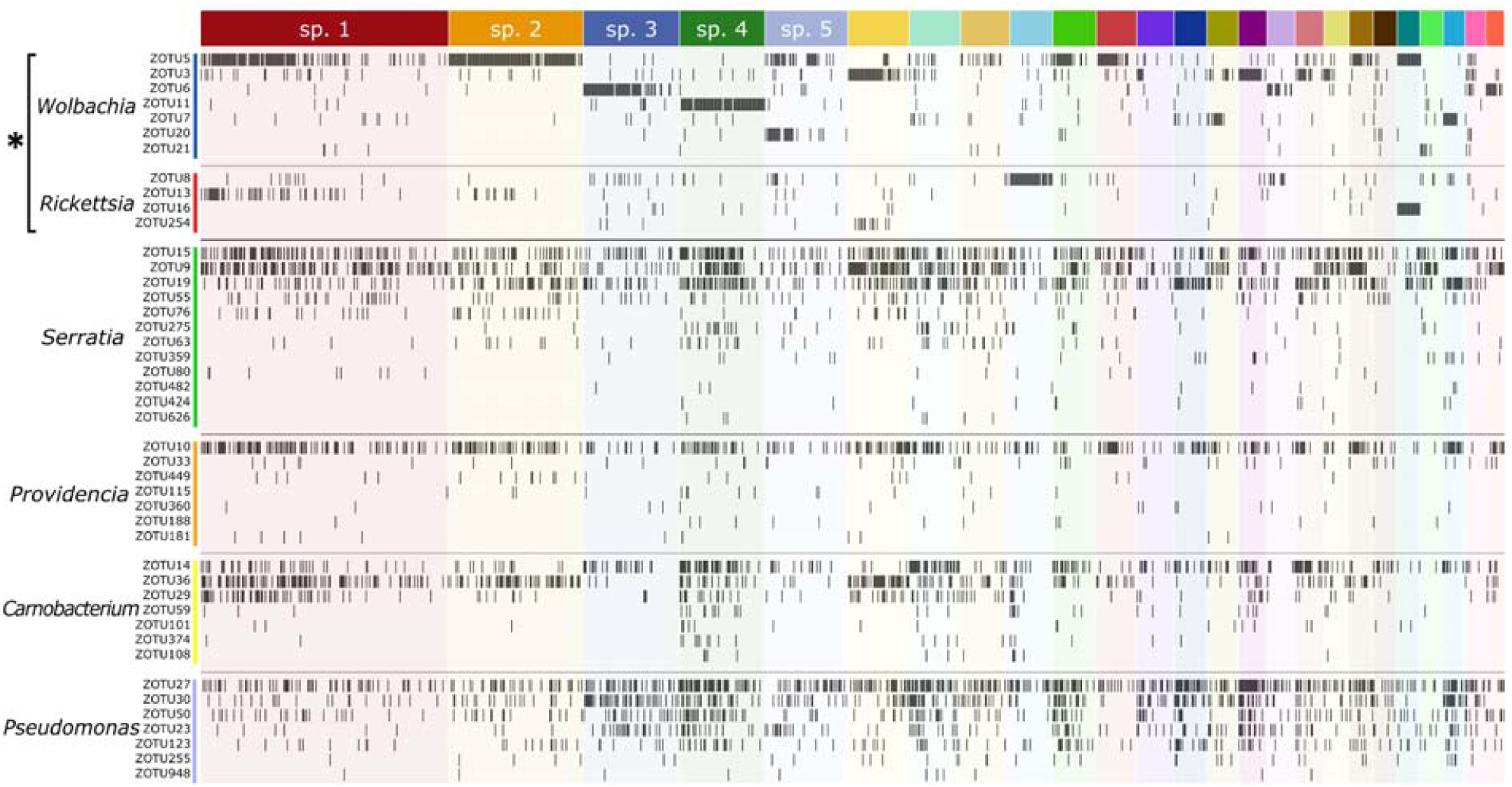
The distributions of dominant bacterial genotypes (ZOTUs) within most abundant OTUs acros host species reveal differences in specificity among bacterial functional categories: facultative endosymbionts (denoted with an asterisk) form relatively specific associations with hosts, unlike other microbes. Insect individuals, represented by columns, are sorted by species, sex, and decreasing bacterial abundance (as in Fig. 2). Rows represent bacterial genotypes (ZOTUs) sorted into 97% OTUs, and black lines indicate symbiont genotypes represented by at least 100 copies of 16S rRNA in specific insect individuals.

Cytochrome oxidase (I) amplicon data for Alphaproteobacteria provide even greater phylogenetic resolution. Specifically, *Wolbachia* comprised most of the 0.5% of non-insect COI sequences in the dataset. Based on these COI data, *Wolbachia* was represented by 2 or more reads and thus scored as present in 862 (47%) of samples, with an average of 269 reads per sample (max 4,875, median 100). These *Wolbachia* COI reads represented 258 unique ZOTUs grouped into 23 97% OTUs. A comparison of the 16S-V4 and COI data revealed > 90% agreement among the two datasets regarding the *Wolbachia* presence in the samples, after applying cutoffs likely to exclude much of likely cross-contamination (3 COI reads vs 400 reads of 16S rRNA) (Supplementary Table S5).

The comparison of *Wolbachia* <u>COI OTU</u> distributions across phorid species (Fig. 5a) resembles *Wolbachia* <u>16S rRNA genotype</u> distribution patterns (Fig. 4). These data show that phorid species typically associate with a single dominant *Wolbachia* COI OTU and differ considerably in infection prevalence. <u>COI genotype</u>-level data for both hosts and symbionts (Fig. 5b) provide much more detailed information: they show that genotypes of the same fly species frequently associate with different genotypes of *Wolbachia*, with substantial variation among genotypes in infection prevalence. Interestingly, we observe some of the same *Wolbachia* COI genotypes across genotypes of different phorid species. For example, *Wolbachia* ZOTU206 infects most genotypes of phorid species 2, at high prevalence, but also colonizes several genotypes of sp. 7, sp. 9, and some less abundant species.

**Fig. 5:**
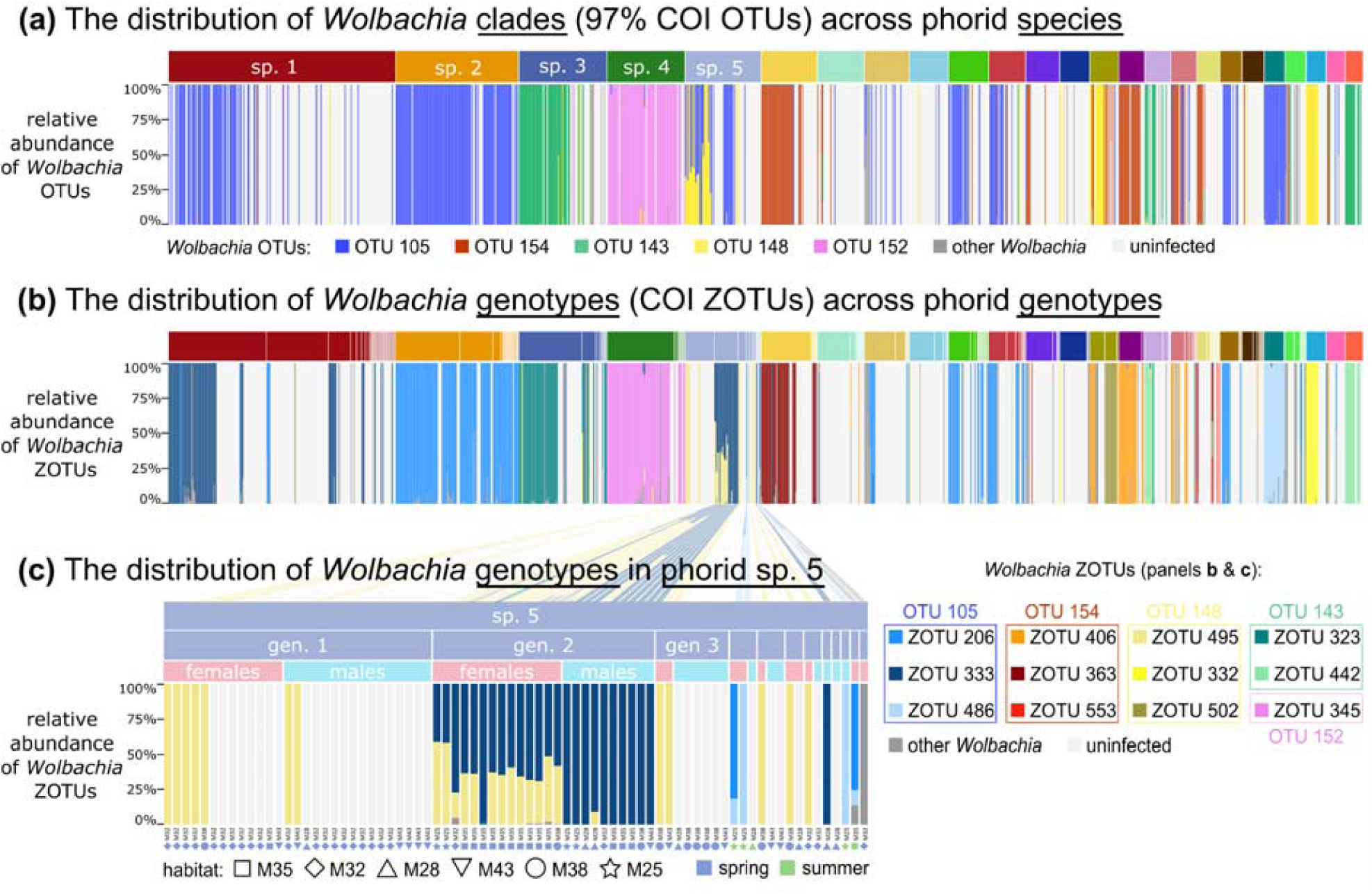
*Wolbachia* COI amplicon data, analysed at different phylogenetic resolution scales, provides much more precise information about the patterns of host-symbiont interactions than 16S rRNA amplicon data. (**a**) The relative abundance of *Wolbachia* COI OTUs among all *Wolbachia* COI reads, across the 25 most abundant scuttle fly species. Individual flies are represented by columns, and sorted by species, sex and decreasing abundance (as in Fig. 2 and 4); (**b**) The relative abundance of *Wolbachia* COI genotypes (ZOTUs) across individual flies sorted into species and additionally COI genotypes; (**c**) The relative abundance of *Wolbachia* genotypes (ZOTUs) among genotypes, sexes, and populations of sp. 5 (*Megaselia* sp. 004), highlights strong specificity in interactions. In all panels, individuals with fewer than 2 *Wolbachia* COI reads are considered uninfected and are represented by white columns.

Within species, we found even greater specificity in host-symbiont interactions concerning sex and site (Fig. 5c). For example, in sp. 5., *Wolbachia* COI ZOTU495 colonised most host genotypes, and tends to achieve higher prevalence in females, only sometimes infecting males. In contrast, *Wolbachia* ZOTU333 infected all individuals - males and females - of one of the abundant phorid COI genotypes, and a single processed male of another genotype, while being absent from others. Interestingly, two other *Wolbachia* genotypes (ZOTU206 & ZOTU486) infected sp. 5 individuals representing different genotypes, all collected in summer rather than spring.

### 3.3. Drivers of microbiota variation

Within our dataset, we observed substantial variation in the prevalence and abundance of microbiota. To quantify the relative contributions of host traits and environmental factors, we turn to our HMSC models of the presence/absence and abundance (conditional on presence) of microbes at three levels of resolution: 16S rRNA OTUs, 16S rRNA ZOTUs, and (for *Wolbachia* only) COI ZOTUs.

#### 3.3.1 Results of the HMSC models

Our predictors (site, season, host species, host sex and host genotype) explained a substantial proportion of the variation in bacterial 16S rRNA OTU and ZOTU presence-absence (Supplementary Table S6), showing high explanatory and predictive power (AUC Δ 0.69). The effects of host individual and site contributed the most to variance, with microbial taxa generally being more prevalent in the spring than summer and more abundant in female than male hosts (Supplementary Table S6). For abundance conditional on presence, the explanatory power remained high, but the predictive power was low due to the strong effect of host individual (Supplementary Table S6). Some individuals had a high abundance of most bacterial taxa, while in others, microbial abundance was consistently low for most bacterial taxa (Supplementary Table S6). Unlike presence-absence, abundance was generally higher in summer than in spring. This indicates that spring samples typically contained many taxa in low abundance, whereas summer samples had fewer taxa in high abundance (Supplementary Table S6).

For *Wolbachia* COI genotypes, presence-absence was strongly shaped by host species and genotype, with additional effects of season and site (Supplementary Table S6, Supplementary Fig. S3c). Model performance was exceptionally high (Supplementary Table S6; AUC Δ 0.90), with most strains being more prevalent in spring and in female hosts. Models of *Wolbachia* abundance conditional on presence showed high explanatory power (AUC = 0.94), but low predictive power – with individual variation driving 36% of the observed differences (Supplementary Table S6). Strain-specific abundances of *Wolbachia* were consistently higher in females than males (Supplementary Table S6).

At the level of individual microbial taxa, explained variance (Tjur’s R^2^) varied significantly, with more prevalent taxa generally showing higher values (Supplementary Fig. S2). The relative contributions of specific drivers also differed among taxa (Fig. 6).

**Fig. 6:**
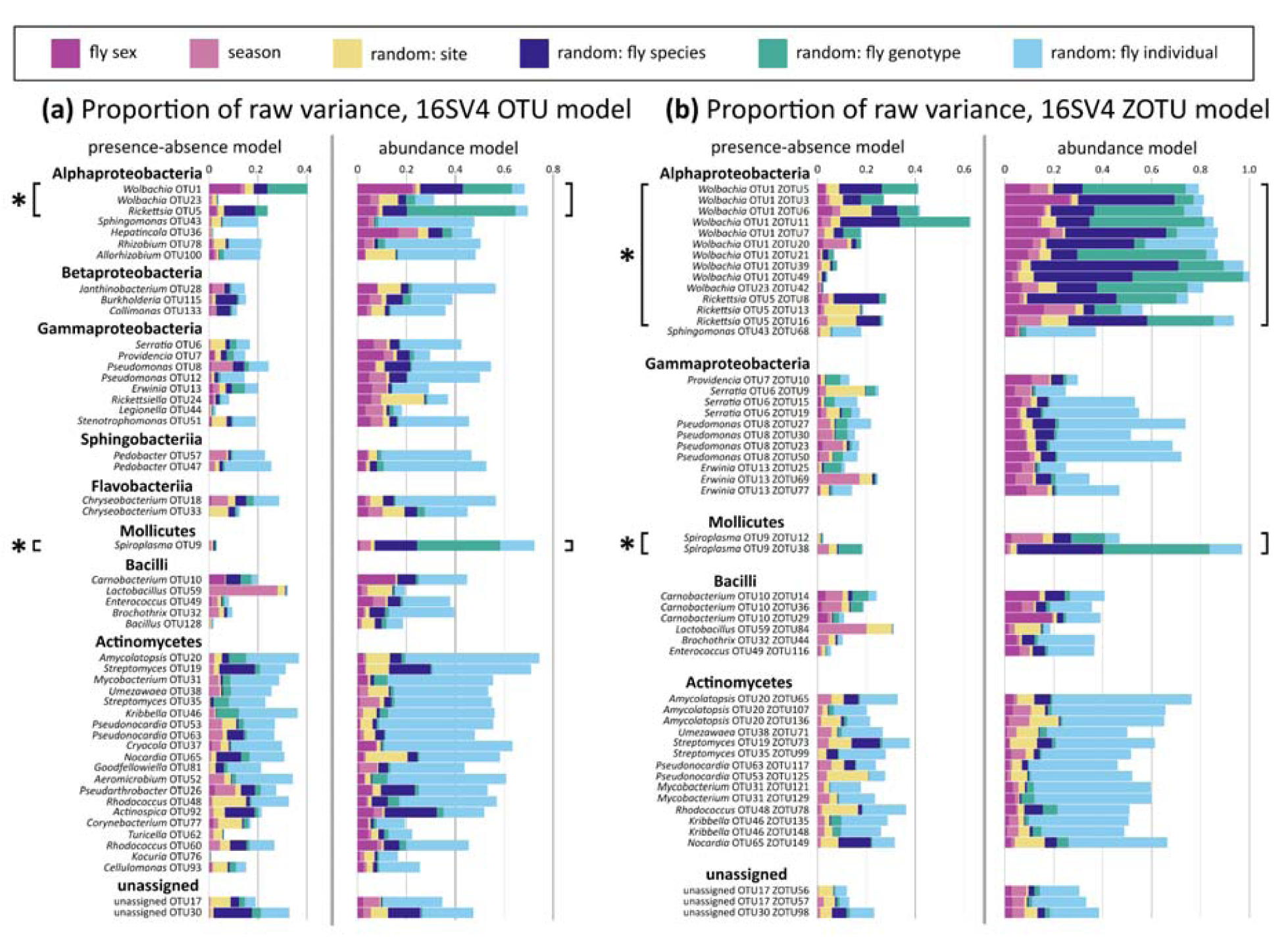
Proportion of variance explained in HMSC models of presence-absence and abundance conditional on presence resolved per microbial (a) 16S-V4 OTU and (b) 16S-V4 ZOTU. Note that compared to other bacteria, facultative endosymbionts (denoted with asterisks) are highly influenced by the host sex, species and genotype. For 16S-V4 ZOTUs, we show a subset of the 50 most abundant ZOTUs (out of 100 ZOTUS included in the models).

**Fig. 7:**
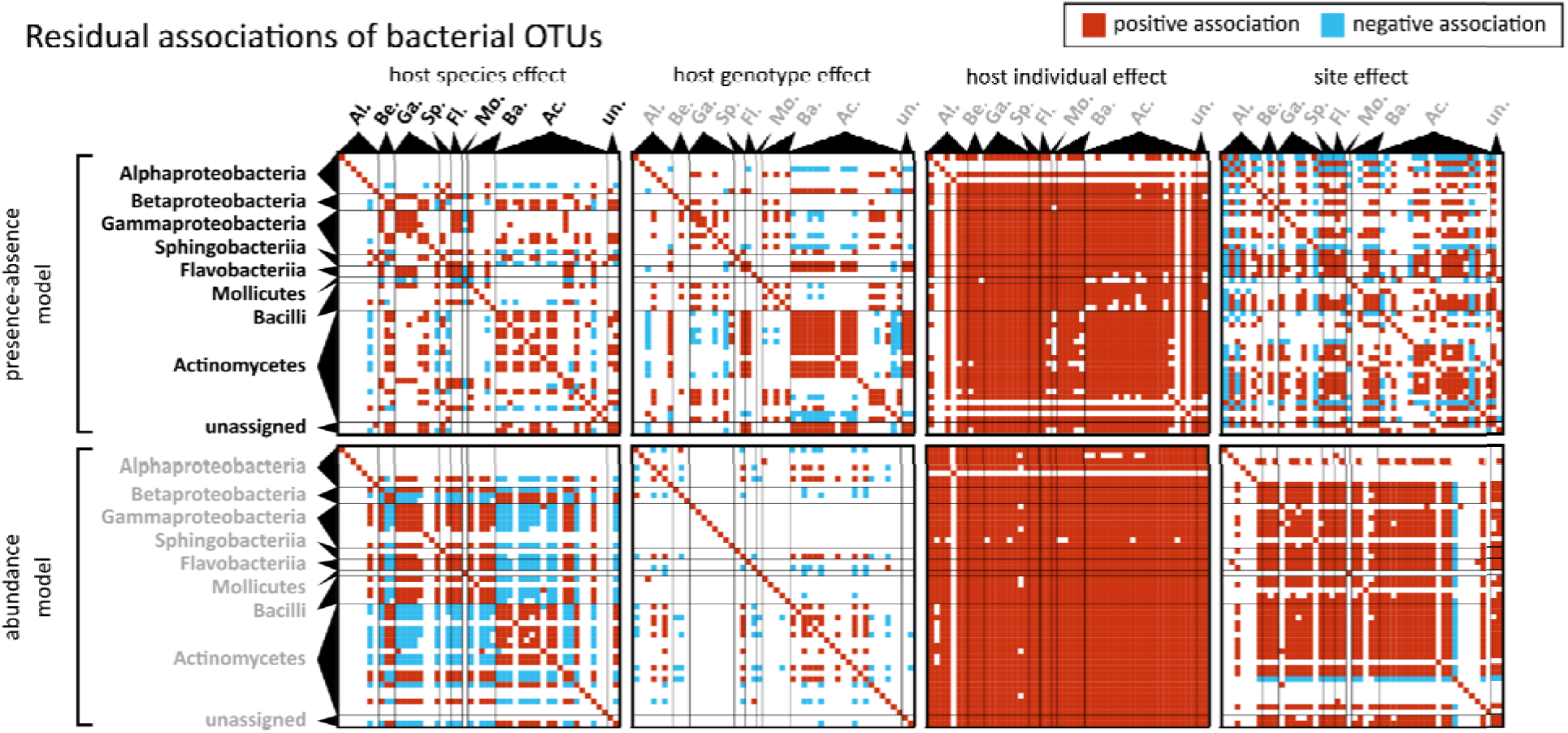
Residual associations among bacterial taxa reveal how OTUs may influence each others’ presence and abundance (conditional on presence) once shared external drivers are accounted for. Th columns of the panels correspond to the four random levels included in the HMSC models, and th rows of the panels represent models of presence-absence (top row) or abundance conditional on presence (bottom row). Within each panel, the bacterial taxa have been sorted into classes (consistent among panels). Red colour indicates pairs of taxa with a positive residual association and blue colour pairs of taxa with a negative residual association, with a posterior probability of 0.90 used as a threshold.

#### 3.3.2 Drivers of variation in bacterial clade occurrence and abundance

The HMSC model explained up to 40% of the variance in the presence-absence of individual 16S-V4 OTUs, with much variation occurring among host individuals (Fig. 6a, Supplementary Fig. S3a). However, for facultative endosymbionts (*Wolbachia*, *Rickettsia*, *Spiroplasma*), the variance was primarily driven by host species, sex, and genotypes, while *Lactobacillus* showed the strongest seasonal variation (Fig. 6a and Supplementary Fig. S3a). For abundance conditional on presence, the model explained up to 70% of the variance, with the lowest contribution from host individual in facultative endosymbionts (Fig. 6a, Supplementary Fig. S3a). Different facultative endosymbionts exhibited distinct patterns: *Wolbachia* varied the most with host species, sex and genotype; *Rickettsia* showed a strong effect of host genotype, and *Spiroplasma* exhibited minimal differences between sexes (Fig. 6a, Supplementary Fig. S3a).

Similar patterns emerged for the 16S-V4 ZOTUs, where models of *Wolbachia* ZOTUs showed the highest Tjur R^2^ values for both presence/absence and abundance for conditional on presence (Supplementary Fig. S3b). For both responses, most variation was attributed to host species and genotype. Among facultative endosymbionts, host individual effects explained most variation (Supplementary Fig. S3b). This was particularly clear in Actinomycetes, which showed the strongest effects of host individual (Supplementary Fig. S3b).

For *Wolbachia* strains identified via COI, variance in presence/absence was primarily shaped by host genotype, species, sex, or site, dependent on the ZOTU (Supplementary Fig. S3c). However, on the overall, a relatively low share of the overall variance was explained by the model (Supplementary Fig. S3c). For abundance conditional on presence, a larger variance proportion wa associated with the host individual, with additional contributions from host species, genotype, and sex (Supplementary Fig. S3c).

#### 3.3.3 Residual associations among microbial taxa

After accounting for the joint impact of systematic effects (3.3.2), we turned to patterns of residual associations. This approach revealed that microbial taxa may influence each other’s presence and abundance *beyond* responding to the same external drivers. Such associations can be addressed at the level of each random factor, i.e. within host species, genotypes, and sites.

At the level of host species, distinct groups of positive bacterial associations were observed. They were strongly related to taxonomy and exhibited positive residual co-occurrence within groups and negative or weak residual co-occurrence between groups. One group was primarily composed of Actinomycetes, while another included Gammaproteobacteria, Bacilli, and Flavobacteria. These patterns were especially evident in OTU abundance conditional on presence. At the level of host genotypes similar clusters emerged. However, in terms of abundance conditional on presence, associations were less consistent and relatively few statistically supported associations in abundance were discovered.

At the level of host individuals, most microbial taxa (except some Alphaproteobacteria and Actinomycetes) showed consistent positive associations among each other, in both presence-absence and abundance conditional on presence. This suggests that host individuals tended to have either very high or very low bacterial diversity, with little structuring among individual microbial taxa. In terms of abundance conditional on presence, most microbial clades were associated positively with each other.

At the level of sites, we found many statistically supported associations. In the presence-absence model, most taxa were positively associated, however, several unrelated OTUs formed negative associations. In terms of abundance conditional on presence, associations were also largely positive, except for some Alphaproteobacteria and Actinomycetes taxa.

Patterns observed for the 16S OTU models were similar to the ones shown for the 16S ZOTU models (Supplementary Fig. S4). In contrast, for the *Wolbachia* models, few significant association were detected, likely due to low statistical power from the limited occurrence of specific ZOTU across host species and individuals (Supplementary Fig. S4).

## 4. Discussion

Insect biodiversity largely consists of poorly known ‘dark taxa’, such as scuttle flies (15,30,59). In these groups, addressing even basic questions, such as about their interactions with other organisms, pose considerable difficulties (29,60). The situation is changing with the advent of new conceptual approaches combined with progress in sequencing, bioinformatics, and statistical modelling (11,17). Our study demonstrates how these innovations, especially when applied in combination, can greatly enhance our understanding of interactions within the diverse wild insect communities.

### 4.1. What have we learnt about scuttle flies?

This study shows how diverse communities of a dark taxon can be accurately resolved *at the same time* as their associations with microbes or other organisms. This approach opens up entirely new prospects for community ecology. To date, most studies of ecological communities have focused on characterising species’ Grinnellian niches – that is, how communities are shaped by the match between abiotic conditions and species requirements (61–63). This has largely neglected how species occurrence is determined by their interactions with other species – the Eltonian niche (64,65). This study characterises both of these niches in one go. By simultaneously identifying the host species (here: scuttle flies) and their associated taxa (here: bacteria) across sites, we may effectively characterise entire wild communities of multi-partite symbiotic entities, or holobionts (66). The current study focuses on a relatively small set of sites, thus constraining analyses of the Grinnellian niche – but another study shows how this can be achieved for broader sample collection (29). By adding the microbial dimension, we show the way forward for the integration of the two (67).

### 4.2. What have we learnt about scuttle fly microbiota?

Our study resolves massive variations in the microbiota of scuttle flies at several levels of organization. We show large variation in bacterial abundance – a rarely assessed, although critical aspect of microbiota (19,20,40). We record up to four orders of magnitude differences in the number of 16S rRNA copies even among individuals representing the same population and sex. This is likely to correlate with large differences in bacterial cell density, although variations in rRNA operon copy numbers within bacterial genomes (68,69), and copies of the genome per cell (70), can also play a role. In turn, bacterial density should correlate with the functional significance of bacteria in the host biology, even if the density-dependence of bacterial effects on the host (71,72) makes the direction of these effects hard to predict.

This variation in bacterial abundance is linked to the community composition. In particular, facultative endosymbionts *Wolbachia* and *Rickettsia* tend to be highly abundant when present, driving the overall abundance among species and sexes. These widespread microbes show highly specific patterns in their distribution, with host species, genotype, and sex explaining a large share of the overall variance across statistical models. Pronounced variation in the association of facultative endosymbiont vs host genotypes suggests that these symbioses are dynamic at relatively short timescales - with symbiont strains moving among host lines, spreading, and likely affecting the host population structure, but rarely persisting for a long time (3,73,74). At the same time, the broad distribution of some *Wolbachia* COI genotypes across different phorid species indicates that they are relatively frequently transmitted horizontally among distantly related host individuals (75).

*Wolbachia* is thought to spread in host populations through the combination of reproductive manipulation and beneficial effects, including protection against natural enemies and nutrient provisioning. Our community-level data strongly suggest that reproductive manipulation is common in the phorid system. Specifically, major differences in *Wolbachia* prevalence and abundance between males and females (Supplementary Table S6) argue for a negative link between infection and male functions. These patterns are consistent with expectations for some types of reproductive manipulation: male killing, feminisation of genetic males, or parthenogenesis induction (3,76). Unfortunately, we lack information about sex ratios in the surveyed populations, since our sampling and specimen selection strategy were not designed to assess sex ratios accurately. Nonetheless, in eight out of the 25 most abundant species sex ratio among processed specimens significantly deviated from an expected 1:1 sex ratio towards females, and in six others towards males (chi-squared test, p < 0.05). The current tentative evidence for reproductive manipulation needs to be systematically validated through the combination of genomics and experiments.

Conversely, many of the remaining abundant bacteria represent generalist clades. For example, the genus *Pseudomonas* is known from a wide range of environments and host organisms that it influences in a multitude of ways. While hundreds of *Pseudomonas* species have been formally named, their environmental variation is much greater (77). The same applies to most other abundant bacterial genera in our dataset, including *Serratia* (78), *Carnobacterium* (79), *Providencia* (80), and others. To date, research on these bacterial genera has focused on a limited set of species of direct relevance to human health and economy, and our understanding of their taxonomic and functional diversity in natural ecosystems has remained limited. At the same time, our 16S rRNA genotype data highlight the surprisingly broad distribution of bacterial genotypes across phorid species, suggesting their recent and likely ongoing transmission among distantly related hosts. Here, the relatively slow rate of evolution in the rRNA gene (81) constrains further resolution of phorid-bacteria relationships. Higher-resolution tools, especially genomics, will be necessary to conclude how these microbes are distributed and move among hosts and environments, and what their effects may be.

Overall, our data suggest that adult scuttle flies may not typically form specific associations with microbes other than heritable facultative endosymbionts. We suspect that at the time of flies’ emergence from pupas, their body surfaces and digestive tracts may be largely sterile - and that only later they become colonised by some of the versatile opportunistic bacteria present in the environment.

### 4.3. What do scuttle flies teach us about broader insect–microbe interaction patterns?

Perhaps the most universal finding of this study, and the key to interpreting most patterns, is the striking contrast between microbial functional categories - facultative endosymbionts, and other bacteria - in their associations with hosts (82). In future studies, it will be critical to distinguish among functional categories when assessing insect-microbe associations. At the same time, when characterising the microbiota of a poorly known host group, distribution patterns can provide insights into microbial biology and its role in the host’s biology.

Facultative endosymbioses, represented here by *Wolbachia*, *Rickettsia*, and *Spiroplasma*, are often thought to form relatively dynamic associations with hosts. We have convincing examples of relatively recent symbiont host switches among related species (74,83) and across distantly related species that form natural communities (84), as well as selective sweeps that they may drive (5,7,85,86). These processes can dramatically influence insect population structure, performance in changing environmental conditions, or interactions with other organisms (11). Conversely, this can affect species’ responses to ecological and environmental challenges and opportunities, their longer-term evolutionary trajectories, as well as broader community processes (4,87). However, we do not know how often these processes take place, or what are their primary drivers and likely consequences (73,75). We believe that the patterns observed in phorids - including a relatively short age of most host-symbiont associations, frequent transmission of some *Wolbachia* lineages across host species, and common reproductive manipulation - will eventually prove to be general features of facultative endosymbiont infections across diverse insects. Pioneering methodological approaches such as those used in our study will enable systematic testing of these hypotheses about facultative endosymbiosis stability and infection dynamics.

In contrast, microbial clades other than facultative endosymbionts are likely to have much broader distributions. Most genera listed in Supplementary Table 4 have been repeatedly reported from many diverse insects from around the world - for example, from butterflies or colonies of army ants from the Neotropics (52,53,88). It is tempting to think of these microbes as highly mobile across global environments and host clades. However, comparative data from vertebrates show how host biology may shape the specificity of bacteria, including strains from broadly distributed genera (89). The systematic assessment of microbial diversity, transmission patterns, and roles across taxa and environments would require not only massive sampling and stringent contamination control (18) but also markers that provide much greater phylogenetic resolution than a short rRNA gene region - ideally, entire genomes.

At the same time, scuttle flies do not represent the full range of insect-microbe associations known. Many other insects form highly stable associations with specific bacterial clades, often maintained through nutritional dependencies: whether with obligate endosymbionts (9) or with host-adapted gut microbiota (82,90). We would normally expect bacteria representing these categories to be always present and abundant within the host species - and in surveys spanning multiple species, to show evidence of co-diversification with hosts (91,92). However, the near-universal associations with *Wolbachia* or *Rickettsia* that some scuttle fly species form may hint towards the indispensable effects of these microbes, as reported in some other insects (93,94).

### 4.4. Advancing the study of wild-insect microbiota: a comprehensive approach

This study introduces a novel methodological framework for understanding wild insect communities and their microbial associations, offering insights far beyond the capabilities of standard marker gene amplicon sequencing, or other methodologies. Our approach incorporates four key innovations: 1: host DNA-based identification done alongside bacterial community characterisation, 2: quantification of bacterial abundance, 3: the use of COI amplicons for high-resolution characterisation of *Wolbachia*, and 4: the application of advanced modelling to decipher complex community patterns of hosts and microbes. By combining these techniques at scale, we provide a nuanced and biologically relevant understanding of microbiota diversity and its ecological role in wild insect communities.

DNA barcoding has already revolutionised the understanding of arthropod biodiversity (95). In this study, we simultaneously characterised microbiota and barcoded their hosts, providing a robust framework for reconstructing insect microbiota patterns at high resolution. As shown in other insect systems, COI data may also enable correcting misidentifications and reconstructing patterns such as parasitoid infections - potentially important confounding variables in studies of microbiota (96).

We incorporated synthetic spike-ins to estimate microbial abundance, a step that adds a crucial dimension to our understanding of microbiota. Such quantification has been repeatedly recommended (17,19,20) and our approach shows how it can be implemented in large-scale studies. While qPCR has been the most popular quantification approach, spike-ins (40) offer a cost-effective alternative for obtaining consistent abundance estimates spanning some six orders of magnitude. The patterns reconstructed are biologically realistic, as revealed by the non-random bacterial distributions observed within scuttle fly species. Quantification helps to assess samples where bacterial abundances are high and community composition reliable, but also identify samples with low bacterial abundances, for which we should treat information on community composition with caution. This is preferable to discarding such samples for being below the detection limit. While this approach awaits more systematic validation, we recommend incorporating spike-in-based quantification into studies of the microbiota across small organisms.

COI amplicon data for *Wolbachia* provide much higher phylogenetic resolution than standard 16S rRNA amplicon data, and thus yield more accurate insights into relationships among insects and these key facultative endosymbionts. This was realised over two decades ago when COI was incorporated into the multi-locus sequence typing (MLST) scheme (97). Interestingly, the realisation that *Wolbachia* co-amplifies with insect mitochondrial COI (98) has motivated dedicated efforts to block its amplification (99). Here we have considered this co-amplification as an opportunity rather than a problem. The > 90% overlap between 16S rRNA-based and COI-based *Wolbachia* detection (depending on the cutoffs used - K. H. Nowak, unpublished) suggests that this symbiont can be reliably detected based on COI data alone. Increased phylogenetic resolution greatly enhances our ability to reconstruct key distribution and transmission patterns. This suggests exciting prospects for extracting infection information from rapidly growing barcoding datasets.

Finally, our implementation of Hierarchical Modelling of Species Communities proved key to dissecting patterns in and drivers of variation among host and microbiota. While HMSC is ideally suited for dealing with the complexities of high-dimensional community data, its applications in the reconstruction of insect-bacteria interaction networks have so far been limited (100). On the other hand, in other host-microbe systems, the method has proven its power to reconstruct complex processes, patterns and their drivers (101). In particular, HMSC offers a solution to disentangling species associations caused by joint preference for the same environmental conditions from interactions among species. Thus, we expect HMSC to serve as an essential statistical framework for understanding broad questions about arthropod-microbe interactions in future large-scale projects.

The solutions outlined above - combined with other recommended approaches such as contamination filtering (17,19,102) - help address the key limitations of amplicon sequencing across large multi-species sample collections (39). Further, the amplicon sequencing approach can be applied in tandem with complementary techniques ranging from high-throughput (meta-) barcoding-based insect surveys on the one end, and high-resolution metagenomics and metatranscriptomics on the other (11,103), to increase both the breadth and depth of the investigation. And while in this study we uncover ecological patterns of fundamental importance in a dark taxon, analogous approaches will help address broader questions about biological interactions within diverse natural communities.

## Acknowledgements

The project was supported by the Polish National Agency for Academic Exchange grant PPN/PPO/2018/1/00015 and Polish National Science Centre grants 2018/31/B/NZ8/01158 and 2021/41/B/NZ8/04526 (to PŁ). EH’s work on Swedish Phoridae was funded by the Swedish Taxonomy Initiative grant 2016-203 4.3. OO and TR were funded by the European Research Council (ERC) under the European Union’s Horizon 2020 research and innovation programme (grant agreement No 856506: ERC-Synergy project LIFEPLAN). OO was additionally funded by the Research Council of Finland (grant no. 336212 and 345110). The authors also acknowledge support from the National Genomics Infrastructure in Stockholm. We thank Dave Karlsson, Esben Bøggild, trap owners, and sample sorters for the Artdatabanken-funded Swedish Insect Inventory Project that provided the samples for this project.

## Supplementary item legends

Supplementary figures, tables and files are available at: https://github.com/KarolHub/Phorid-Microbiota/tree/main/Supplementary_materials

**Supplementary Fig. S1:**
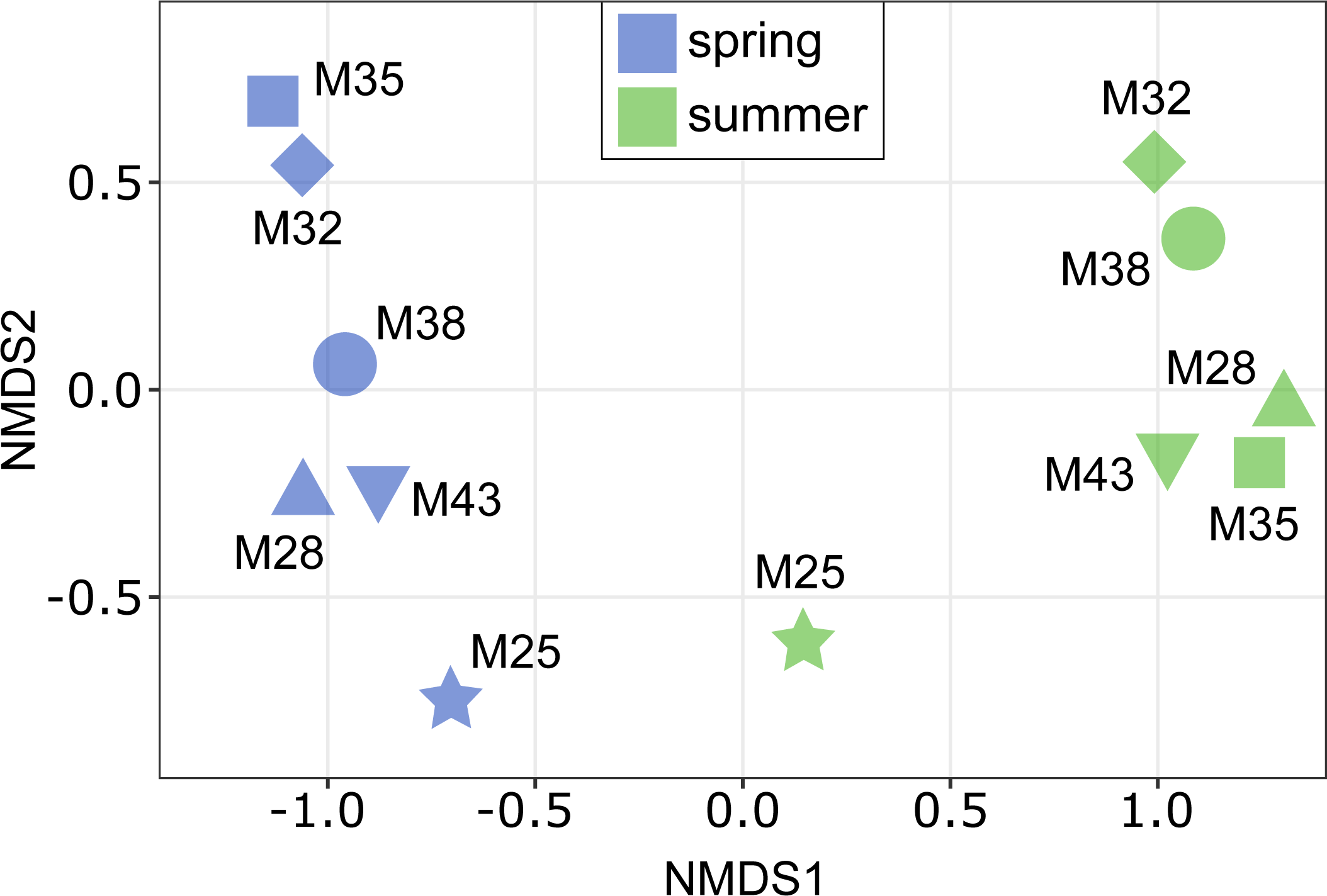
Ordination of host communities composition among sites and seasons (NMDS plot based on Bray-Curtis dissimilarity).

**Supplementary Fig. S2:**
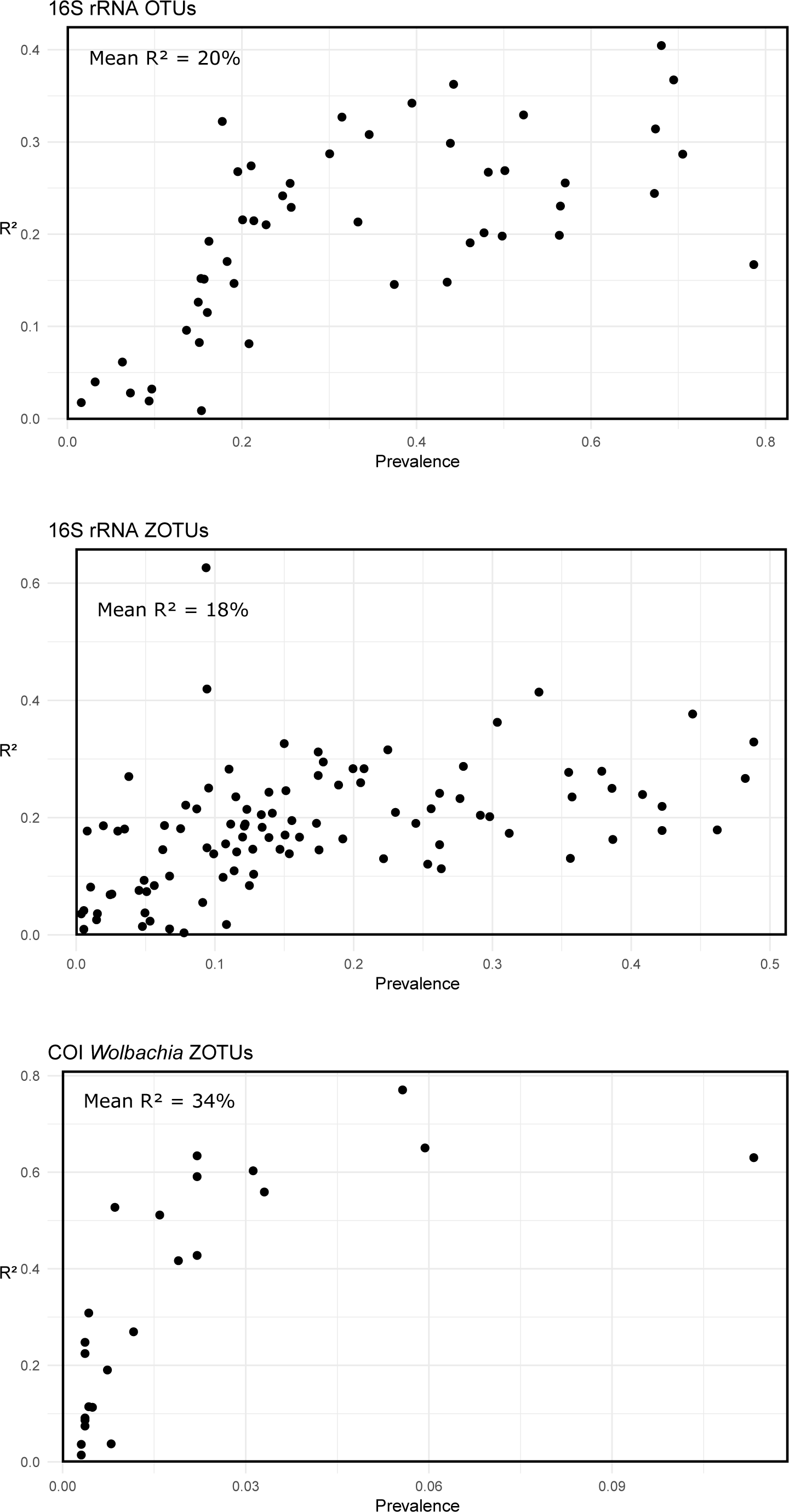
Relationship between the explained variance (R^2^) of bacterial taxa presence and their prevalence.

**Supplementary Fig. S3:**
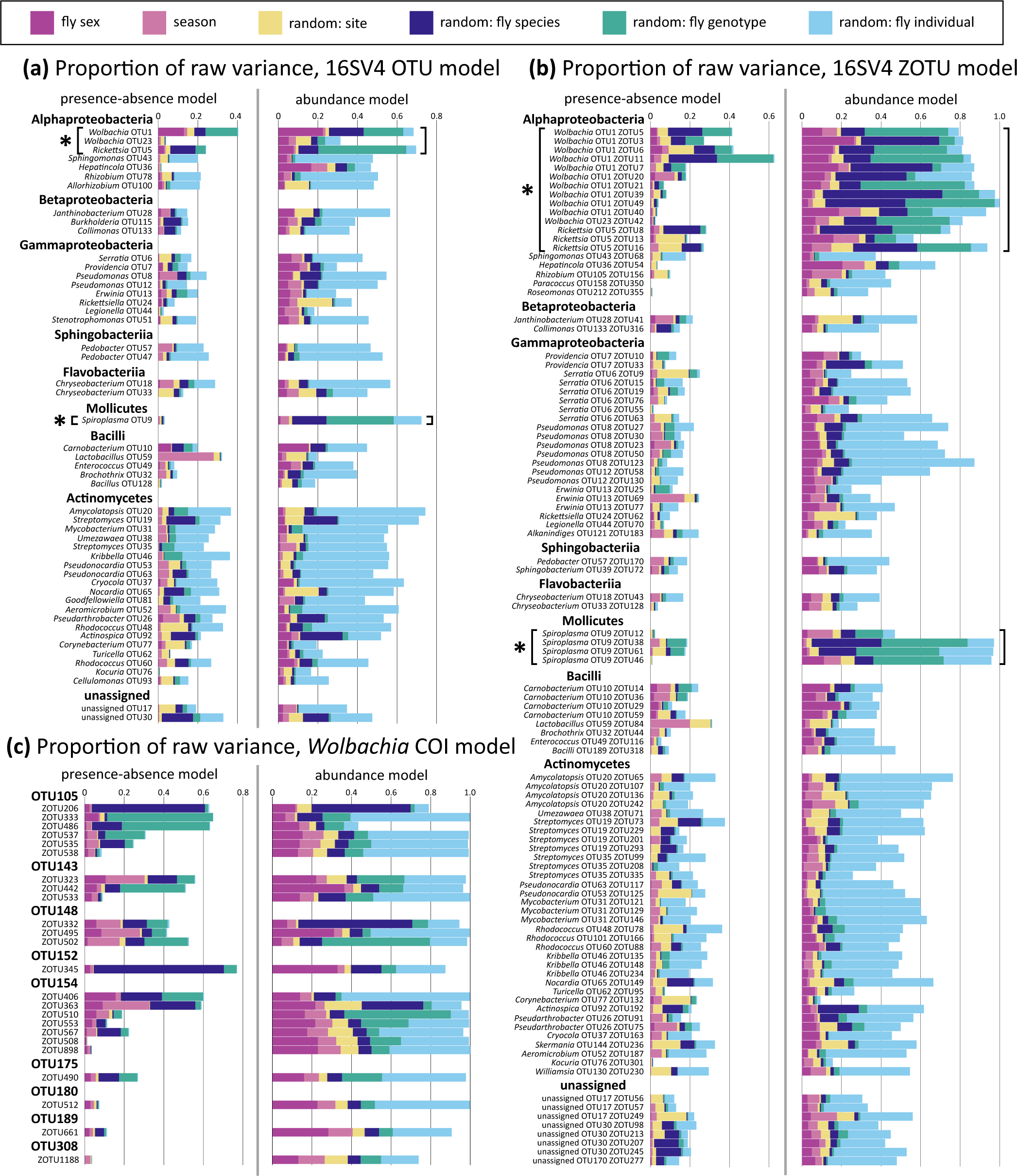
Proportion of variance explained per microbial taxon by the presence-absence model and the abundance conditional of presence model at the levels of microbial 16S-V4 OTUs (a), 16S-V4 ZOTUs (b) and *Wolbachia* COI ZOTUs (c).

**Supplementary Fig. S4:**
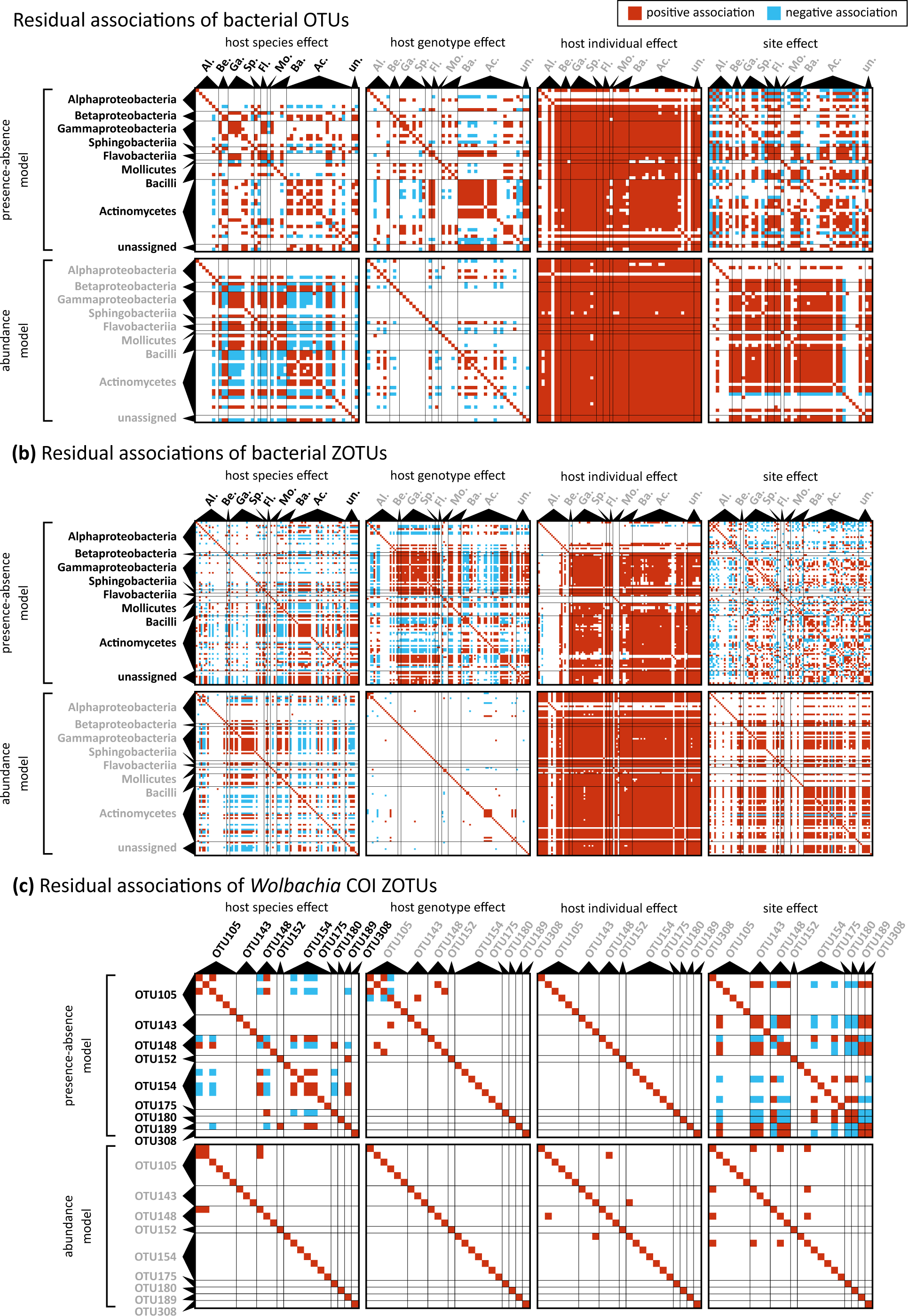
Residual associations in the presence-absence model and the abundance conditional of presence model among bacterial taxa at the levels of microbial 16S-V4 OTUs (a), 16S-V4 ZOTUs (b) and *Wolbachia* COI ZOTUs (c).

Supplementary Table S1: Details of the sampling sites and the numbers of processed insect specimens passing the quality criteria.

Supplementary Table S2: List of sample barcodes, species and genotype assignments, along with their sexes, sampling locations and accession numbers.

Supplementary Table S3: Numbers and proportions of 16S rRNA reads classified as contaminants, spikeins, and symbionts across the samples, identified by the QUACK script.

Supplementary Table S4: List of 16S rRNA OTUs, their prevalence and abundance across samples.

Supplementary Table S5: Comparison of *Wolbachia* presence between COI and 16S-V4 datasets.

Supplementary Table S6: Results of the HMSC models

Supplementary File 1: Compressed table of COI ZOTUs read numbers and taxonomic assignments across the samples.

Supplementary File 2: Compressed, decontaminated table of 16S-V4 ZOTU read numbers and taxonomic assignments across the samples.

Supplementary File 3: Compressed, decontaminated table of estimated 16S-V4 ZOTU copies and taxonomic assignments across the samples.

